# Adaptive encoding speed in working memory

**DOI:** 10.1101/2022.09.08.507070

**Authors:** Joost de Jong, Hedderik van Rijn, Elkan Akyurek

## Abstract

Humans can adapt when complex patterns unfold at a faster or slower pace, for instance when remembering a grocery list that is dictated at an increasingly fast pace. Integrating information over such timescales crucially depends on working memory, but although recent findings have shown that working memory capacity can be flexibly adapted, such adaptations have not yet been demonstrated for encoding speed. In a series of experiments, we found that young adults encoded at a faster rate when they were adapted to overall and recent rate of incoming information. Interestingly, our participants were unable to use explicit cues to speed up encoding, even though these cues were objectively more informative than statistical information. Our findings suggest that adaptive tuning of encoding speed in working memory is a fundamental but largely implicit mechanism underlying our ability to keep up with the pace of our surroundings.

**Significance Statement:** Humans can store information very quickly. For instance, when we hear someone speak twice as fast as normal, we can still follow quite well. How is this possible? We hypothesized that when humans expect limited time to store a piece of information (e.g. when listening to a sped-up podcast) they would ideally store that information more quickly before it’s gone. Indeed, we found that young adults encoded more information per second when they implicitly expect that they have little time to do so. However, they were unable to use explicit cues about how much time they have. It seems that young adults can, at least implicitly, tune the pace at which they store information.

## Introduction

Whether it is following an increasingly rapid story told by an excited friend or listening to a piece of music that slows down its tempo; humans are experts at keeping up with the pace of our surroundings. For instance, humans rather easily comprehend speech at two-fold increases in speech rate (Foulke & Sticht, 1969). But what underlies our ability to track information efficiently over such a wide range of timescales? A common bottleneck in domains like music and speech is working memory (Schulze & Koelsch, 2012). Ideally, information should be encoded sufficiently before the stimulus is no longer there, especially when interfering stimuli may follow. Intuitively, if we expect information to arrive at a fast rate, information should also be encoded at a faster rate.

Despite this intuitive idea that we can adapt working memory encoding speed to the expected stimulus duration, such adaptations have not been tested empirically. This hiatus is surprising for three reasons. First, adaptive dynamics have been demonstrated in a wide variety of domains, such as sensory adaptation (Fairhall et al., 2001; Wark et al., 2009), decision-making (Murphy et al., 2021; Ossmy et al., 2013), timing (Remington et al., 2018), event integration (Akyürek et al., 2008), speech perception (Lerner et al., 2014), learning (Behrens et al., 2007; Piray & Daw, 2020), and motor adaptation (Gonzalez Castro et al., 2014). Overall, these lines of research show that the speed of cognitive (or neural) processes adjusts to the timescale of (or the rate of change within) the environment. For instance, the rate of evidence accumulation in decision-making speeds up when incoming evidence changes more frequently (Glaze et al., 2015).

Second, from its very inception (e.g., Sperling, 1963), working memory research has had a strong focus on encoding speed (see also, Bays et al., 2011; Bundesen, 1990; Busey & Loftus, 1994; Gegenfurtner & Sperling, 1993). However, while many factors have been found to modulate encoding speed, such as luminance/contrast (Loftus & Ruthruff, 1994), attention (Reinitz, 1990), and working memory load (Bays et al., 2011; Vogel et al., 2006), none of these qualifies as an adaptation to the temporal structure of the environment. One possible exception is the finding that encoding speed is modulated by the overall foreperiod preceding stimulus onset (Vangkilde et al., 2012, 2013). However, these adaptations are with respect to properties that facilitate preparation for memoranda, not the temporal properties of the memoranda themselves.

Third, adaptive processing in working memory has been observed with regard to its capacity. State-of-the-art computational models assume that capacity is adaptively allocated to memoranda (e.g., van den Berg & Ma, 2018). Indeed, it has been shown empirically that working memory capacity adapts to stimulus statistics (Orhan et al., 2014), can be flexibly allocated to information that is likely to be relevant (Bays & Husain, 2008) and can be controlled at will to match task demands (Machizawa et al., 2012). However, none of these accounts have considered that encoding speed may adapt to the rate at which information arrives at the senses.

In this paper, we study whether and how working memory encoding speed, in a similar way as capacity, is adapted to the structure of the environment. Specifically, we hypothesize that encoding speeds up when the expected stimulus duration is brief. In a series of experiments, we leverage distribution effects, serial dependence, and cueing to systematically manipulate expectations about stimulus duration. To preview our results, we found that humans encode information about twice as fast when they information to be presented briefly, but only when these expectations were induced implicitly. These findings suggest that implicit encoding speed adaptations may underlie our ability to keep up with the pace of our surroundings.

## Materials and Methods

### Participants

Participants were first-year psychology students (mean age 20.9 years; 63% female) at the University of Groningen who obtained partial course credit for participating. Informed consent was obtained before the experiment started. Participants with an average capacity estimate of higher than 50 rad^-1^ (see Analysis) were excluded from further analysis (four in experiment 1, one in experiment 2, four in experiment 3). Indeed, their average capacity far exceeded the Q3 + 3*IQR criterion for outlier detection, which was 8.0 (experiment 1), 9.3 (experiment 2) and 10.9 (experiment 3). Using this stricter exclusion criterion did not change the interpretation of our results. For experiments with bootstrapping (experiment 1 and 3), our original exclusion criterion (capacity above 50 rad^-1^) was applied to each bootstrap. As a result, we discarded a participant within a bootstrap 5% and 15% of the time in experiment 1 and experiment 3, respectively.

### Apparatus and stimuli

The experiment was programmed in OpenSesame, a free software tool for designing experiments (Mathôt et al., 2012). Stimuli were presented on a 19 inch CRT screen with a resolution of 1,280 × 1,024 pixels, running at 100 Hz. Participants were seated in a sound-attenuated room with dimmed lights approximately 60 cm from the screen. A gray background was maintained during the entire experiment. Memory items were Gabor patches with a gaussian envelope (spatial frequency: 0.05 cycles/degree, std: 12 pixels, phase: 0), presented at fixation. Its orientation was picked from a uniform distribution spanning 0 to 180 degrees. Memory items were masked with 50 overlapping Gabor patches with random orientations (spatial frequency: 0.05 cycles/degree, std: 10 pixels, phase: 0) and their centers scattered in a 40×40 pixel square.

### Procedure

Participants performed several practice trials in a delayed estimation task (Wilken & Ma, 2004). A trial proceeded as follows. A fixation dot was shown for 500 to 1000ms (uniformly sampled), after which the memory item was presented at fixation. The presentation time (i.e., for how long the memory item was physically on the screen) was 50, 200 or 400ms. To control the time available for encoding, we attempted to eliminate any retinal or cortical after-image by presenting a pattern mask for 100ms immediately after memory item offset. Participants had to retain the memory item over the next 1000ms, after which a ‘wheel probe’ was presented. This was a Gabor patch that participants could turn around by moving their mouse to the remembered orientation and clicking to confirm their response. Participants only received trial-to-trial feedback in practice trials. In the real experiment, only block-wise feedback (total number of points per block) was presented. Apart from the counterbalancing of presentation times, trials per block and the inclusion of a cue at the start of each trial (Experiment 3), the procedure was identical across experiments.

### Experiment 1: Blocked presentation time

In our first experiment (N=57), we tested whether a higher overall presentation increases encoding speed in working memory. To this end, each block either had predominantly brief presentation time (50ms: 70%, 200ms: 15%, 400ms: 15%), or long presentation time (50ms: 15%, 200ms: 15%, 400ms: 70%). Each presentation time regularity lasted for four blocks of 40 trials (i.e., 160 trials in total), after which participants switched (from brief to long presentation time, or vice-versa). Participants completed a total of 640 trials and nine practice trials at the start of the experiment. Participants were not informed about overall presentation time within a block or switches in overall presentation time between blocks. Because the size of encoding speed adaptations is unknown, we could not base our sample size on formal power calculations. Instead, we sampled batches of participants until we obtained strong evidence for or against our hypothesis with our Bayes Factor (i.e. BF_10_ > 10 or BF_10_ < 0.1). We further ensured that each condition contained enough trials to have reliable estimates of the circular standard deviation of responses (see, Bays et al., 2011).

### Experiment 2: Sequential presentation time

In this experiment (N=24), we assessed whether encoding speed would also adapt to recent, sequential changes in presentation time. We counterbalanced the order of presentation times with a de Bruijn sequence, using the software described in (Aguirre et al., 2011). A de Bruijn sequence perfectly counterbalances the order of its elements up to a certain degree using the smallest sequence length. We designed the sequence such that each possible order of three presentation times (e.g., 50 - 50 - 400) would occur with equal frequency within each block. This allowed us to fit an encoding curve for each n-1 and n-2 presentation, using an equal number of trials for each presentation time on trial n. Participants performed a total of 675 trials and nine practice trials at the start of the experiment. Sample size was determined in the same way as Experiment 1.

### Experiment 3: Cued presentation time

In our final experiment (N=19), we explored whether adaptations in encoding speed could be induced with explicit cues that are highly informative. At the start of each trial, we presented either the word ‘FAST’ in red, or ‘SLOW’ in blue for 400ms. These cues were 80% valid. ‘Fast’ cues predicted 50ms presentation times 80% of the times (with 10% for both 200 and 400ms), and vice versa for ‘slow’ cues. Participants were informed that the cues were informative and encouraged to make use of the cues. They completed a total of 500 trials and 20 practice trials at the start of the experiment. Sample size was determined in the same way as Experiment 1. However, contrary to our expectations, we found that ‘fast’ cues decrease encoding speed after collecting the first batch of participants. This constituted evidence against our one-sided hypothesis (i.e., that ‘fast’ cues increase encoding speed), but our Bayes factor tests the presence of an effect, not its direction. Therefore, to see whether we were justified to stop data collection, we estimated the one-sided Bayes Factor (BF_>_) using the approximation in Morey & Wagenmakers (2014) and obtained strong evidence against our hypothesis that cues increase encoding speed (BF_10>_ = 0.098).

### Analysis

All analysis was performed in R (R Core Team, 2018; version 3.5.1). Error angles were computed as the reproduced angle minus the actual angle of the memory item multiplied by two (since Gabors have a rotational symmetry of 180 degrees instead of 360 degrees), such that errors spanned -180 to 180 degrees. We quantified memory precision as the inverse of the circular standard deviation (σ^-1^) of the error angles at each presentation time, t. We subtracted the memory precision that would be expected for that sample size (N), which was computed by taking inverse of the mean of 1,000 circular standard deviation estimates from circular uniform distributions with sample size N.

To quantify encoding speed and capacity in working memory, we fitted exponential encoding curves (see e.g., Bays et al., 2011; Busey & Loftus, 1994) for each participant in each condition with two parameters: Maximum capacity (c) and encoding speed (τ^-1^):

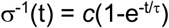

These parameters map onto the asymptotic memory precision for very long presentation times (c, see Fig. 1), and encoding speed (i.e., the speed at which precision reaches that asymptote; (τ^-1^, see Fig. 1). Effects on maximum capacity (c) become most apparent for long presentation times because stimuli are remembered better (or worse) even when they have been presented for a very long time. In contrast, effects on encoding speed are most apparent for brief durations, because two encoding curves may reach the same asymptote but at different speeds. To illustrate, a fast encoding curve will reach a high precision early on, compared to a slow encoding curve (see Figure 1, bottom right panel). Nevertheless, the slow encoding curve eventually catches up with the fast encoding curve, reaching the same asymptote. These different behavioral signatures of capacity and encoding speed effects allow us to reliably separate them when fitting encoding curves. It is important to note that all types of trials in Experiment 1 and Experiment 3 (including invalidly cued and ‘block-inconsistent’ trials) were used to fit encoding curves, because it is impossible to estimate capacity and encoding speed using only a single presentation time. Encoding curves were fitted with the nls.multstart package (Padfield & Matheson, 2020; version 1.2.0), which initializes multiple starting values for each parameter when fitting the encoding curve and selects the set of parameters with the lowest AIC score (in our case, the highest likelihood, because the number of parameters remained constant). We normalized the parameters by dividing estimates by the mean estimate for each participant, which allowed us to assess changes in capacity or encoding speed as induced by our experimental manipulations.

**Fig. 1.**
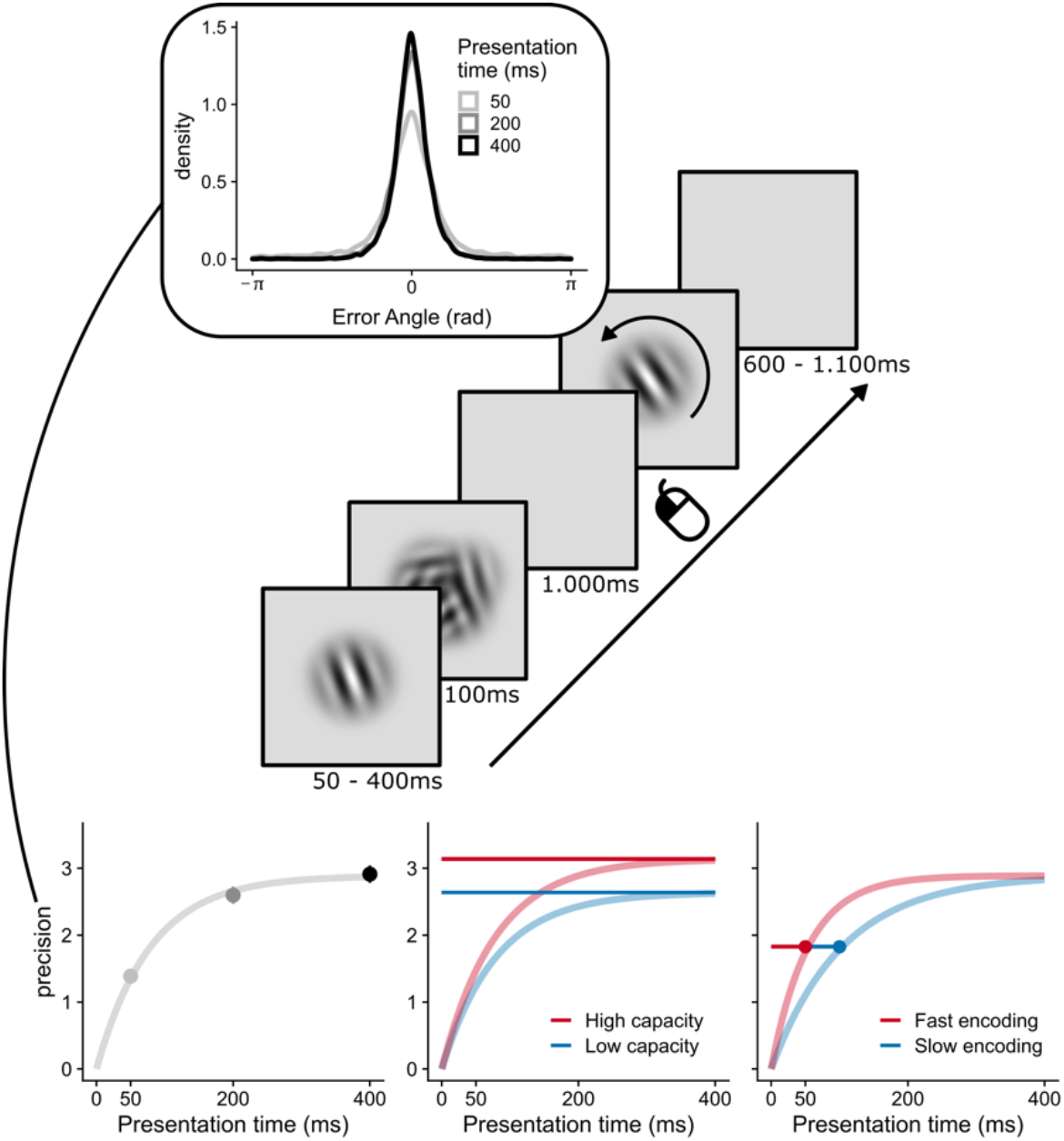
Experiment and analysis setup. Top: experiment setup. Participants had to remember the orientation of a Gabor that is presented for 50, 200 or 400ms and immediately followed by a pattern mask. After 1.000ms, participants had to adjust the orientation of a second Gabor to reproduce the contents of their memory. Inset: the distribution of errors (i.e., reproduced angle minus target angle). Bottom: We computed the inverse of the circular standard deviation of those errors to characterize memory precision, which smoothly increased with presentation time. An exponential encoding curve is fitted (the leftmost panel), which has two parameters: Maximum capacity and encoding speed. The middle panel expresses differential effects of capacity (high in red, low in blue) with fixed encoding speed. While both curves reach a different asymptote, they reach that asymptote equally fast. The right panel depicts the effects of encoding speed (fast in red, slow in blue) with fixed capacity. Both curves reach the same asymptote, but they reach the same precision at different points in time.

The effect of expected presentation time on normalized encoding speed was assessed by fitting a linear regression model. The null model predicted normalized encoding speed with only normalized capacity as a predictor. The alternative model added the manipulation of expected presentation time. The same rationale was followed when testing for effects on normalized capacity: The null model only contained normalized encoding speed and the alternative model included expected presentation time. For experiment 1, blocked presentation time was dummy coded (1 for ‘short’ block, 0 for ‘long’ block). For experiment 2, n-1 and n-2 presentation time were added as continuous predictors. For experiment 3, cued presentation time was dummy coded (1 for ‘short’ cue, 0 for ‘long’ cue). To quantify evidence for a modulation of encoding speed by expected presentation time, we performed model comparison using an approximation to Bayes Factors (Wagenmakers, 2007):

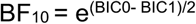

where BIC0 is the Bayesian-Information Criterion (BIC) score for the null-model only containing normalized capacity as a predictor and BIC1 is the BIC score for the model that also includes expected presentation time. High values for BF_10_ reflect evidence against the null and therefore suggest an effect of expected presentation time on encoding speed.

Experiment 1 and 3 were not balanced due to the nature of the experimental manipulation. That is, to induce expectations about presentation time, some presentation times contained more observations than others in certain conditions. Estimates of circular standard deviation are biased substantially at low sample sizes, especially at high standard deviation (see Supporting Fig. S4). Such bias could have a complex impact on the estimated capacity and encoding speed. Therefore, we bootstrapped the entire analysis for those experiments by subsampling (without replacement) for each participant and each condition to the lowest number of observations minus one. We sampled without replacement because estimates of circular standard deviation have higher bias and variance when duplicates can be sampled (see Supporting Fig. S5). We reported the median of all test statistics (regression coefficients, t-values, p-values, Bayes Factors and differences in BIC scores). For plotting, we computed the standard error (SE) of the bootstrap by dividing the estimated standard deviation (SD) by ΔN for each condition. All experimental materials, data and analysis code are available on the Open Science Framework (https://osf.io/d5rnu/?view_only=d4a46b8b900a4c1d8be69b9b57481cc3).

## Results

### Encoding speed adapts to overall presentation time

In our first experiment, we tested whether humans could increase their encoding speed when the overall presentation time was short. Humans are sensitive to the overall distribution of temporal intervals that they perceive (e.g., Jazayeri & Shadlen, 2010). Therefore, if participants tune their expectations to the overall stimulus duration, they should also tune their expectations about overall presentation time and adapt their encoding speed accordingly. To test this hypothesis, we varied the overall presentation time between blocks. Participants alternated between blocks of 160 trials where presentation time was either short (50ms) or long (400ms) for 70% of the trials. We hypothesized that, if humans adapt to overall presentation time, they should have a higher encoding rate in the blocks with brief presentation times compared to those with long presentation times. In line with our hypothesis, encoding speed was systematically faster in blocks with predominantly brief presentation times (*β* = 0.149, t = 3.76, p < 0.001, BF_10_ = 89.82, ΔBIC = 9.00; Fig. 2). We also found that blocks with brief presentation times increased capacity (*β* = 0.09, t = 3.29, p = 0.001, BF_10_ = 19.11, ΔBIC = 5.90). Note that adaptations in encoding speed were independent from increases in capacity, because we controlled for normalized capacity in our analysis. These results suggest that our participants did adapt their encoding rate to the overall presentation time.

**Fig. 2.**
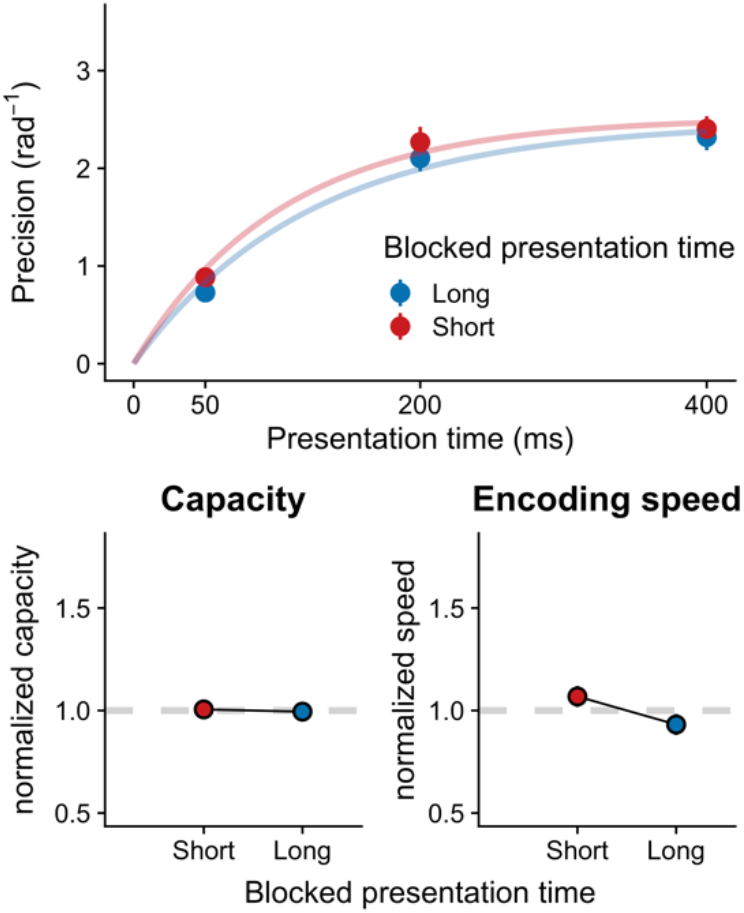
Experiment 1: The effect of blocked presentation time on working memory capacity and encoding speed. The top figure shows how memory precision (mean ± SE) increases with presentation time, which is accurately captured by encoding curves. Blocks either had brief presentation time (red; 50ms presentation time, 70%) or long presentation time (blue; 400ms presentation time, 70%). The bottom figure shows how normalized parameter values (mean ± SE, with an overall mean of 1) vary with rate. The left panel represents capacity and the right panel represents encoding speed.

### Encoding speed adapts to recent presentation time

In our second experiment, we tested how quickly humans can adapt their encoding rate on a trial-to-trial basis. To this end, we leveraged serial dependence observed in the perception of durations. Humans can optimally track stimulus duration on a trial-to-trial basis (de Jong et al., 2021; Glasauer & Shi, 2021). For instance, when a previous stimulus was presented briefly, the next stimulus is also expected to be relatively brief. Therefore, if the previous presentation time is brief, participants should increase their encoding speed in anticipation of the next brief stimulus. In line with our hypothesis, we found that the shorter the presentation time on the previous trial, the faster the encoding speed on the current trial (Fig. 3; *β* = -0.00053, t = -2.68, p = 0.009, BF_10_ = 4.21, ΔBIC = 2.88). This finding suggests that humans speed up encoding when they expect that they have relatively little time for doing so. In contrast, we found evidence against an effect of the previous presentation time on capacity (*β* = -0.00006, t = -0.616, p = 0.54, BF_10_ = 0.14, ΔBIC = -3.88), suggesting that participants specifically tuned their encoding speed to expected presentation time, not capacity. In line with continued adaptation, we also found faster encoding when the previous two stimuli had the same presentation time (Fig. 4; *β* = -0.00178, t = -5.08, p < 0.001, BF_10_ > 10,000, ΔBIC = 18.49), but again no effect for capacity (*β* = -0.00029, t = -0.897, p = 0.373, BF_10_ = 0.18, ΔBIC = -3.45). Notably, a comparison of the regression coefficients (Clogg et al., 1995) showed that the effect of the previous two stimuli on encoding speed was stronger than the effect of the most recent stimulus alone (Z = -4.686, p < 0.001).

**Fig. 3.**
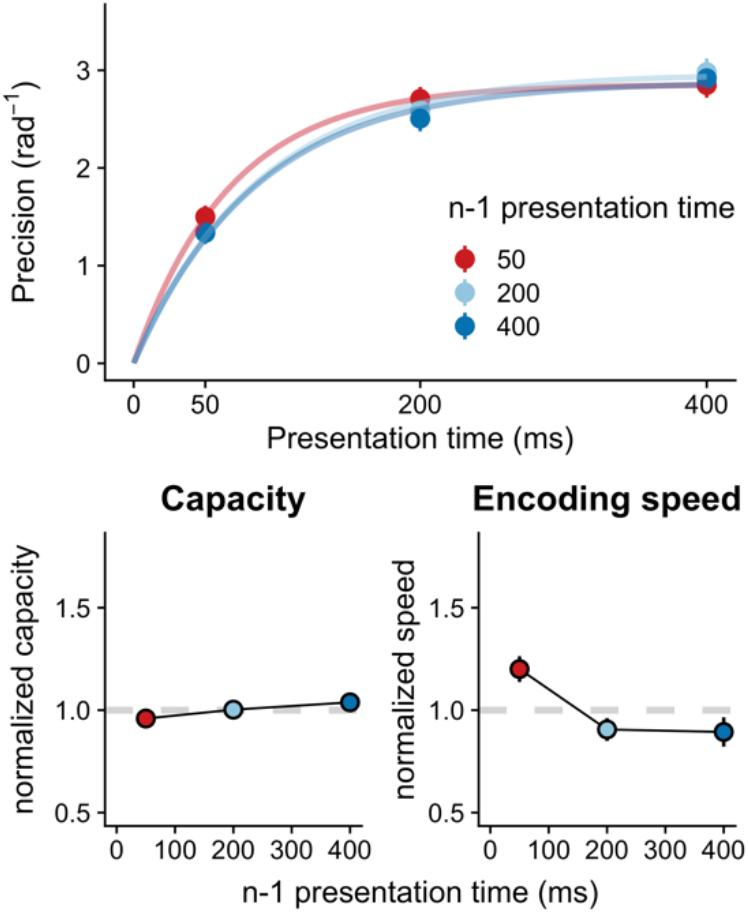
Experiment 2: The effect of n-1 presentation time on working memory capacity and encoding speed. The same plotting conventions are used as in Fig. 2. Colors indicate whether the previous trial had a long presentation time (blue), or brief presentation time (red).

**Fig. 4.**
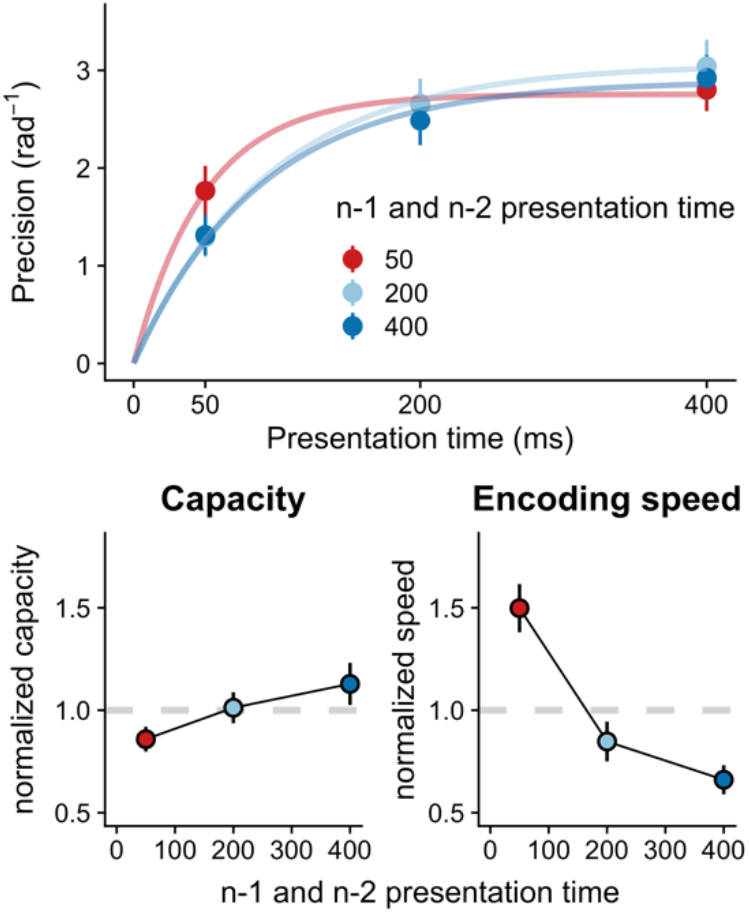
Experiment 2: The effect of n-1 and n-2 presentation time on working memory capacity and encoding speed. The same plotting conventions are used as in Fig. 2. Colors indicate whether the previous two trials both had a long presentation time (blue), or brief presentation time (red).

Our hypothesis predicts that encoding speed specifically adapts to expected presentation time, and not temporal variables that are unrelated to the memory items. Indeed, we found evidence that adaptation was specific to presentation time of the stimulus, not the rate at which the task proceeds after a response. The previous interval between fixation onset and stimulus onset did not significantly modulate encoding speed (*β* = -0.095, t = -1.37, p = 0.175, BF_10_ = 0.23, ΔBIC = - 2.98; Supporting Fig. S1) or interact with the effect of previous presentation time (*β* = -0.001, t = - 0.29, p = 0.774, BF_10_ = 0.09, ΔBIC = -4.80). In the same linear regression model, the effect of previous presentation time was relatively consistent (*β* = -0.0006, t = -2.542, p = 0.012, BF_10_ = 2.24, ΔBIC = 1.61). This finding may seem at odds with reports in the literature that encoding speed in visual detection is modulated by expected foreperiod (Vangkilde et al., 2012, 2013). However, unlike these studies, we did not manipulate foreperiod in a block-wise manner and not over a broad temporal scale (0.3 - 10s). Hence, while expected foreperiod could have some effect on encoding speed, it is unable to account for encoding speed adaptations specific to presentation time. Overall, our findings suggest that humans specifically track presentation time on a trial-to-trial basis and adapt their encoding speed accordingly.

An alternative explanation for our adaptive encoding findings is that participants adjusted their encoding speed to correct for errors on previous trials (similarly to post-error improvements of performance, Danielmeier & Ullsperger, 2011). When large errors were made on previous trials, participants may have tried to compensate by increasing their encoding speed. Hence, this account would predict a faster encoding speed following trials with large errors. However, when we did a median split on the magnitude of the errors, error magnitude on previous trials did not modulate encoding speed on current trials. When participants made large errors on previous trials, their encoding speed on subsequent trials was the same, or if anything slightly lower (*β* = - 0.121, t = -1.69, p = 0.094, BF_10_ = 0.374, ΔBIC = -1.98; Supporting Fig. S2). In sum, it seems that adaptations in encoding speed cannot be explained by error-related corrections.

### Encoding speed is insensitive to explicit cues about presentation time

So far, we have shown that encoding speed is adapted to expectations about presentation time induced by distribution- and sequential-like effects. However, it is unclear whether these expectations can also be induced explicitly, like with spatial cueing (e.g., Jonides, 1981). We ran an experiment where, at the start of each trial, participants were cued (‘FAST’ or ‘SLOW’) whether the upcoming stimulus would be brief (50ms) or long (400ms) with 80% cue validity. Crucially, even though these cues were objectively more informative than our previous sequential and block-wise manipulations, they did not modulate encoding speed. If anything, ‘fast’ cues resulted in a decrease in encoding speed (Fig. 5; *β* = -0.064, t = -0.77, p = 0.321, BF_10_ = 0.306, ΔBIC = -2.36). Indeed, a one-sided Bayes Factor (Morey & Wagenmakers, 2014) suggests that ‘fast’ cues do not increase encoding speed (BF_10>_ = 0.098). We also found no effect of cue on capacity (*β* = 0.052, t = 0. 957, p = 0.19, BF_10_ = 0.47, ΔBIC = -1.52). These findings are in line with several reports in the literature showing that explicit cues are relatively ineffective in guiding expectations about temporal features (Los et al., 2021; Maaß et al., 2019). Indeed, some have shown that temporal expectations are efficiently triggered by implicit associative cues (Salet et al., 2022). In sum, the ineffectiveness of highly informative cues suggests that adaptation to presentation time is more easily triggered by implicit processes, than by explicit cues.

**Fig. 5.**
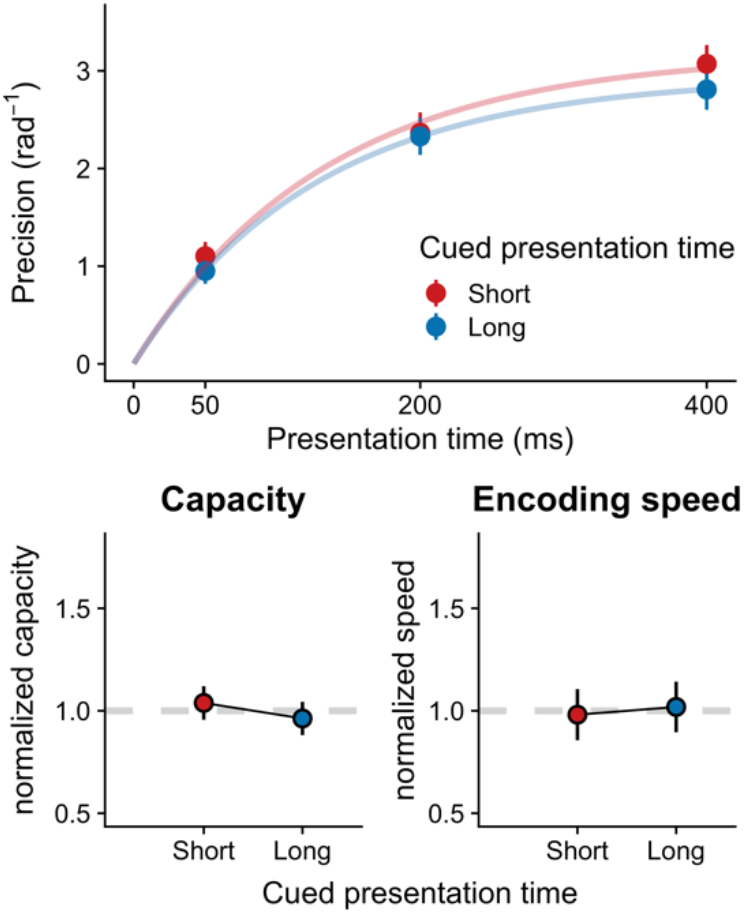
Experiment 3: The effect of cued presentation time on working memory capacity and encoding speed. Cues either predicted brief or long presentation times (50 vs 400ms, 80% validity). The same plotting conventions are used as for Fig. 2.

### Exploring a ‘post-encoding’ explanation

An alternative explanation for our effects shifts their locus to processes that take place after encoding. For instance, when a brief presentation time is expected, participants could more effectively suppress the ensuing mask when it appears early in time. Similarly, when a long presentation time is expected, participants could more easily suppress the mask when it appears later in time. In other words, performance is best when the expected presentation time matches the actual presentation time. This account successfully explains why performance is mainly enhanced at brief presentation times when they are expected. Unlike our theory, however, it also predicts a relative benefit for long presentation times whenever they are expected. Another unique prediction is that when intermediate presentation times are expected, performance should be best at those presentation times.

We systematically tested the ‘post-encoding’ explanation across all our experiments and found that it captures some features of the data that are not easily explained by our adaptive encoding speed theory (Supporting Figs 6-9). For instance, in Experiment 2, we observe some relative costs for long durations when brief ones are expected, whereas our theory predicts no differences (Supporting Fig 7-8). Nevertheless, the alternative explanation is unable to account for a lack of costs at longer presentation times (Experiment 1; Supporting Fig 6) and its unique prediction for intermediate presentation times could not be confirmed (Supporting Fig 7-8). Also, the alternative explanation did not reproduce the asymmetry in our findings, namely that effects of expected presentation time were largest at brief presentation times and gradually tapered off. Even though these behavioral analyses suggest a limited role for post-encoding processes, it would be worthwhile to track both encoding and ‘post-encoding’ processes as they unfold in real time (e.g., through electrophysiological recordings). In sum, these analyses suggest that post-encoding processes cannot solely account for our data, but rather that adaptations to presentation time - at least partly - manifest during working memory encoding.

## Discussion

How do we keep up with the rate at which information reaches our senses? We hypothesized that a fundamental underlying mechanism may be the adaptation of working memory encoding speed to expected presentation time. Indeed, we found that encoding speed was adapted to overall presentation time. Moreover, encoding speed continually and specifically adapted to recent presentation time on a trial-to-trial basis. We further showed that encoding speed adaptations could not be induced by highly informative cues, suggesting that these adaptations may be largely implicit. Overall, our findings suggest that the speed of working memory encoding is optimally adapted to the timescale of incoming information.

Our hypothesis was partly inspired by adaptive dynamics that are found in a wide range of perceptual, cognitive and motor domains. Indeed, we demonstrate that encoding speed in working memory may show similar adaptation to the timescale of incoming information. But what is the underlying logic behind adapting the speed of cognitive processes to match the timescale of the environment? State-of-the-art theories of adaptive dynamics assume that task-relevant variables are tracked in an optimal way. These models propose that, in addition to tracking the mean value of some task-relevant variable, humans also track how quickly that variable is changing (Glaze et al., 2015; Ossmy et al., 2013; Piray & Daw, 2020; Wark et al., 2009). As an example (see Ossmy et al., 2013), when trying to spot a faint signal in the distance, we do not only take into account its luminance, but also its duration. In other words, we keep track of how quickly luminance is changing. If we expect a brief signal, we quickly accumulate evidence before the signal is absent. However, this comes at the expense of accumulating noise. Therefore, if we expect a long signal, we accumulate evidence at a slower rate to prevent false alarms. Indeed, in a decision-making task, Ossmy et al., (2013) found that the rate of evidence accumulation is adapted to the overall duration of the to-be-detected signal. We have shown an analogous encoding speed adaptation in working memory. Given these commonalities, it seems that these formal models of adaptive dynamics (e.g., Glaze et al., 2015) could be a powerful way to conceptualize encoding speed adaptations in working memory.

Another possible idea is that, while increments in encoding speed may improve performance, they incur a commensurate increase in some behavioral or neural cost (for a similar ‘resource-rational’ account of working memory capacity, see van den Berg & Ma, 2018). Whatever form these costs may take, a natural assumption is that humans attempt to maximize their net gain (i.e., performance minus ‘encoding costs’). The second assumption is that participants weigh performance at certain presentation times by their subjective probability. That is, if participants believe that very brief presentation times are more likely (e.g., when the previous duration is short or in the ‘short’ blocks), they put more weight on performance at brief presentation times. These simple assumptions would naturally predict that increasing the rate of encoding is beneficial when brief presentation times are expected.

Given that humans can speed up working memory encoding, how might the brain achieve such a speedup? The analysis of our behavioral data provides a natural clue: The encoding curve we fit to our data (Equation 1) describes the behavior of a leaky integrator that is given a step-like input. Here, the amplitude of the input corresponds to the maximum capacity, while the speed of integration relates to encoding speed. Since we observed independent modulations in encoding speed, the speed of neural integration would be responsible for our effects. However, a plausible alternative is that amplified inputs do not only result in a higher measured capacity, but if capacity has a limit, some of these effects also show up in the encoding rate. To put it simply, a stronger input will drive memory precision to the limit at a faster rate without resulting in much change in that limit. Then, we would expect that increases in encoding speed would be accompanied by little to no increase in capacity. However, we found the opposite: Normalized encoding speed generally had a negative effect on normalized capacity (Supporting Fig. S3). That is, when encoding speed increases, capacity generally decreases. Indeed, this suggests that our results may be explained by a speedup in how quickly populations of neurons change their overall firing pattern (Sohn et al., 2019) or perhaps a speedup in biophysical processes in individual neurons (Durstewitz, 2003), not necessarily changes in input amplitude.

A related question pertains to the locus of encoding speed adaptations. Can our findings be explained by faster temporal summation in the visual system (Loftus & Ruthruff, 1994), as opposed to adaptations in working memory per se? For instance, one might wonder if our results could be explained by contrast adaptation: If the previous stimulus is presented for only 50ms, its contrast could be perceived as lower because temporal summation is not finished yet. Then, on the next trial, adaptation to this lower contrast speeds up temporal summation in the visual system. However, adaptation to low contrast often entails slower changes in firing rates in the visual system, not faster changes (Lesica et al., 2007). Similarly, if temporal summation is completed around ∼100ms (Gorea, 2015), we might still be able to observe differences in encoding when comparing n-1 and n-2 for 200 and 400ms. However, we did not observe significant adaptation around this longer timescale (t = 1.42, p = 0.17). It may nevertheless still be possible that the timescale of adaptation is related to overall encoding speed. When encoding speed is relatively fast to begin with, any increase in encoding speed mainly affects performance at brief presentation times. Therefore, manipulating overall encoding speed (e.g., through contrast or attention) may uncover encoding speed adaptations beyond timescales typically associated with temporal summation in the visual system.

This work constitutes a first demonstration that working memory encoding speed adapts to the rate of incoming information. However, it remains to be seen whether these results generalize to more complex features, different modalities or alternative working memory tasks. Similarly, future work needs to establish whether our results generalize to populations other than the one sampled here, such as children, older adults or non-human animals. Also, while we have argued that the locus of these adaptations is to be found in speedups of neural processes, these claims should be tested using neurophysiological measures with high temporal resolution (e.g. EEG or MEG). Lastly, while we believe that encoding speed adaptations are more easily elicited by implicit cues, it remains to be seen whether participants can explicitly report on the presentation time and whether this explains individual differences in adaptation magnitude.

In conclusion, humans implicitly speed up encoding in working memory when they expect information to be briefly available. These adaptations may be understood in terms of optimally tracking changes in the environment by adapting the speed at which those changes are encoded. Further, additional analyses suggest that encoding speed adaptations are probably not due to amplified visual inputs but may instead constitute actual speedups in neural processes themselves. We believe that encoding speed adaptations in working memory are critical for optimal performance in environments whose pace may change suddenly and substantially.

## Acknowledgments

We would like to thank Thies Hoffman for helping with data collection and two anonymous reviewers for their constructive comments.

## Notes

http://www.osf.io/d5rnu/

### Competing Interest Statement

The authors have declared no competing interest.

### Summary of Updates

This version of the manuscript has added clarifications on encoding curves, describes the results in terms of previous presentation times (instead of information rate) and includes additional analyses concerning alternative explanations for our data.

