## Supplementary Materials for "Adaptive encoding speed in working memory"

**Supporting Information for ‘Adaptive Encoding Speed in Working Memory’**

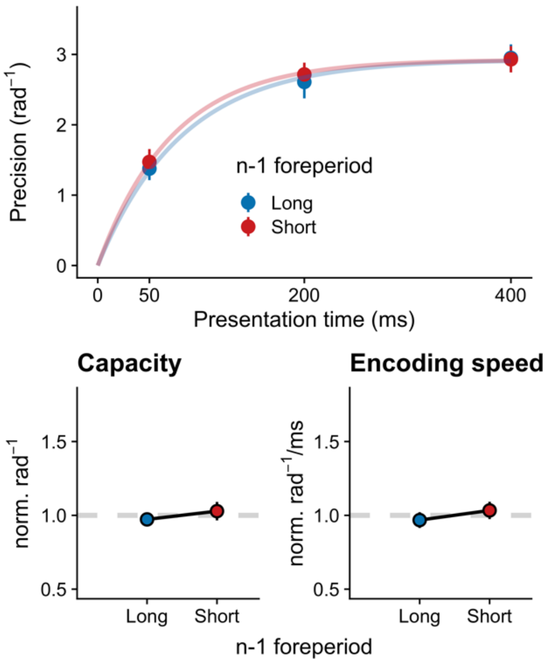


**Supporting Fig. S1.** No effect of the previous foreperiod on working memory capacity and encoding speed. We performed a median split on the previous foreperiod (long in blue; short in red). The same plotting conventions are used as for Figure 2.

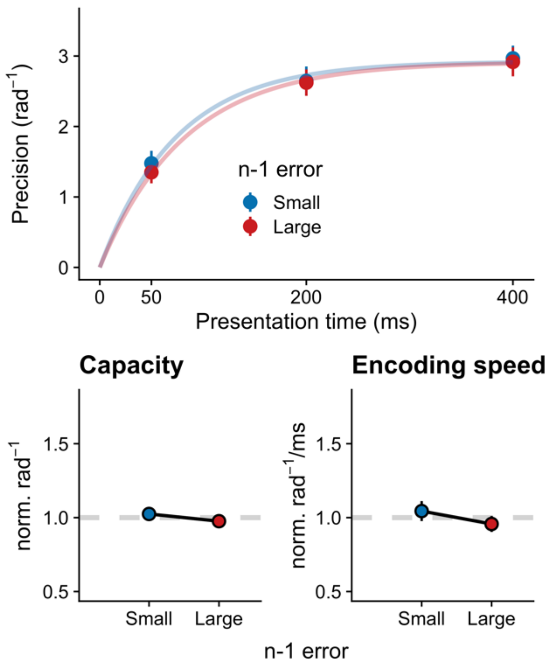


**Supporting Fig. S2.** No effect of the previous error on working memory capacity and encoding speed. We performed a median split on the previous error (small in blue; large in red). The same plotting conventions are used as for Figure 2.

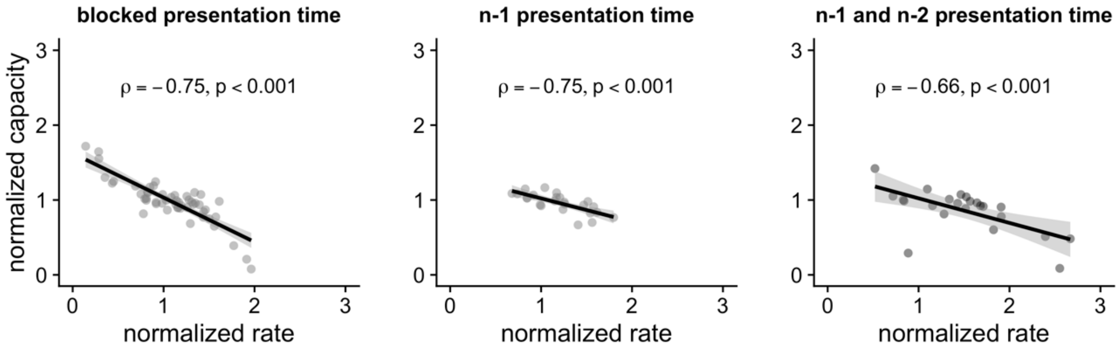


**Supporting Fig. S3.** The effect of normalized rate on normalized capacity for manipulations for which we found adaptive encoding speed. Spearman's rho and associated p-value is reported for each analysis.

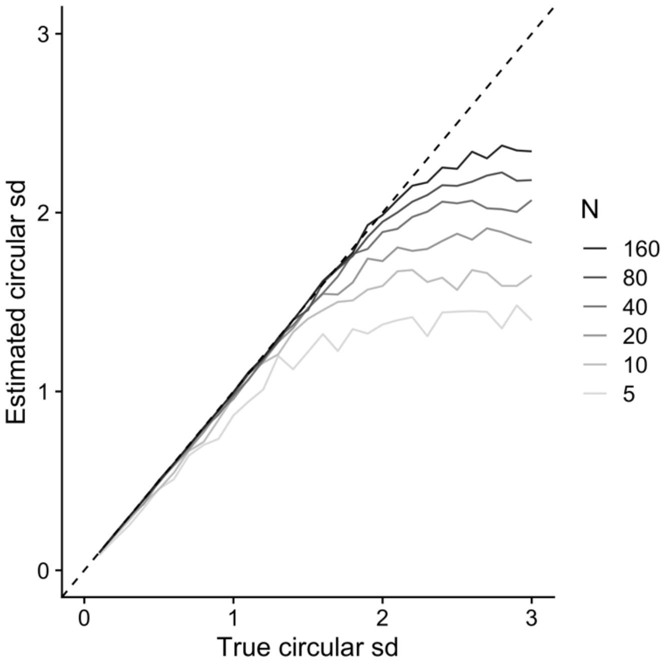


**Supporting Fig. S4.** Estimates of circular sd are biased substantially at low sample sizes, especially for low sd.

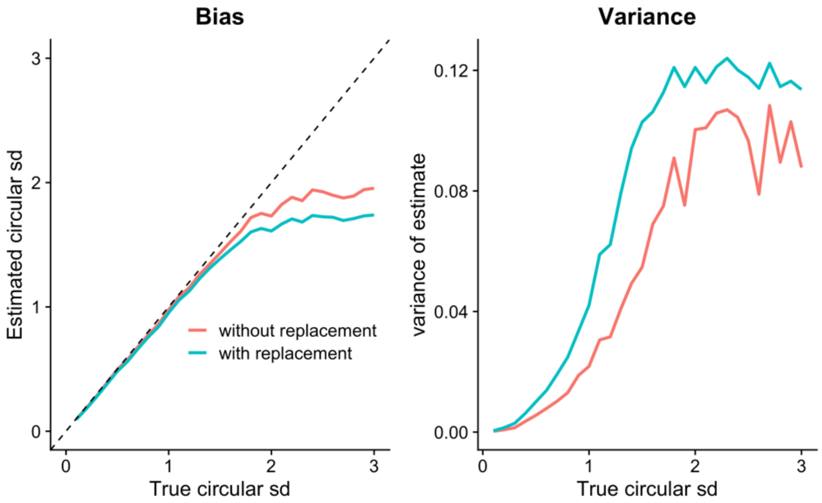


**Supporting Fig. S5.** Bootstrapped estimates of circular sd are biased substantially when sampling with replacement as compared to sampling without replacement, especially for low sd. The variance of the estimates is higher for sampling with replacement. In these simulations, sample size was 25, which is typical for our experiments.

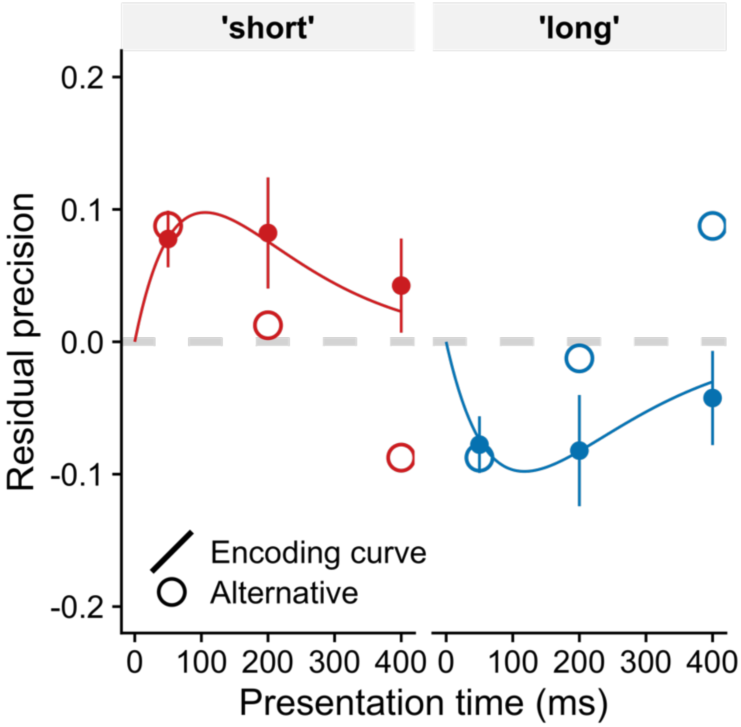


**Supporting Fig. S6.** Testing the alternative explanation that performance benefits and costs scale linearly with the ‘unexpectedness’ of the presentation time (i.e., absolute difference between actual and expected presentation time). Experiment 1: Residual precision (precision – grand average) is plotted at each presentation time for short and long blocks. Filled dots with error bars represent data, smooth encoding curve represents encoding speed adaptations and open circles represent alternative explanation. Encoding speed adaptations were fitted for each condition while keeping capacity fixed. The magnitude of benefits and costs were fitted for each condition separately.

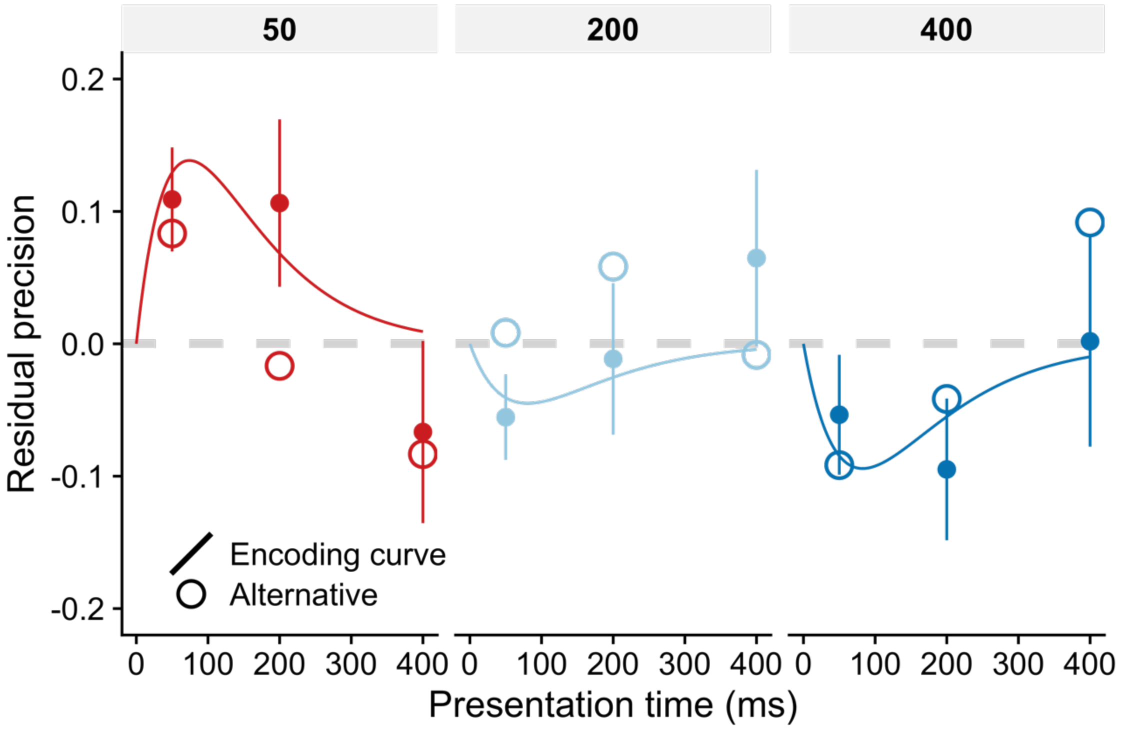


**Supporting Fig. S7.** Testing the alternative explanation that performance benefits and costs scale linearly with the ‘unexpectedness’ of the presentation time (i.e., absolute difference between actual and expected presentation time). Experiment 2 (n-1 analysis): Residual precision (precision – grand average) is plotted at each presentation time for short and long n-1 presentation times. Filled dots with error bars represent data, smooth encoding curve represents encoding speed adaptations and open circles represent alternative explanation. Encoding speed adaptations were fitted for each condition while keeping capacity fixed. The magnitude of benefits and costs were fitted for each condition separately.

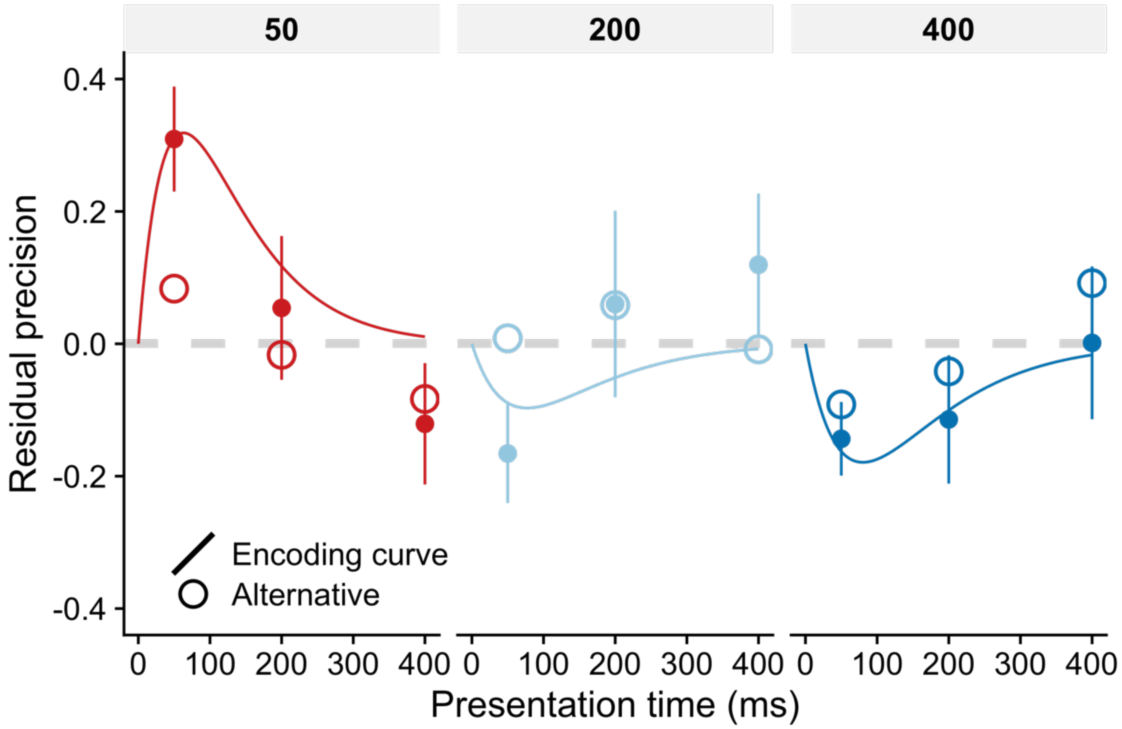


**Supporting Fig. S8.** Testing the alternative explanation that performance benefits and costs scale linearly with the ‘unexpectedness’ of the presentation time (i.e., absolute difference between actual and expected presentation time). Experiment 2 (n-1 and n-2 analysis): Residual precision (precision – grand average) is plotted at each presentation time for short and long n-1 and n-2 presentation times. Filled dots with error bars represent data, smooth encoding curve represents encoding speed adaptations and open circles represent alternative explanation. Encoding speed adaptations were fitted for each condition while keeping capacity fixed. The magnitude of benefits and costs were fitted for each condition separately.

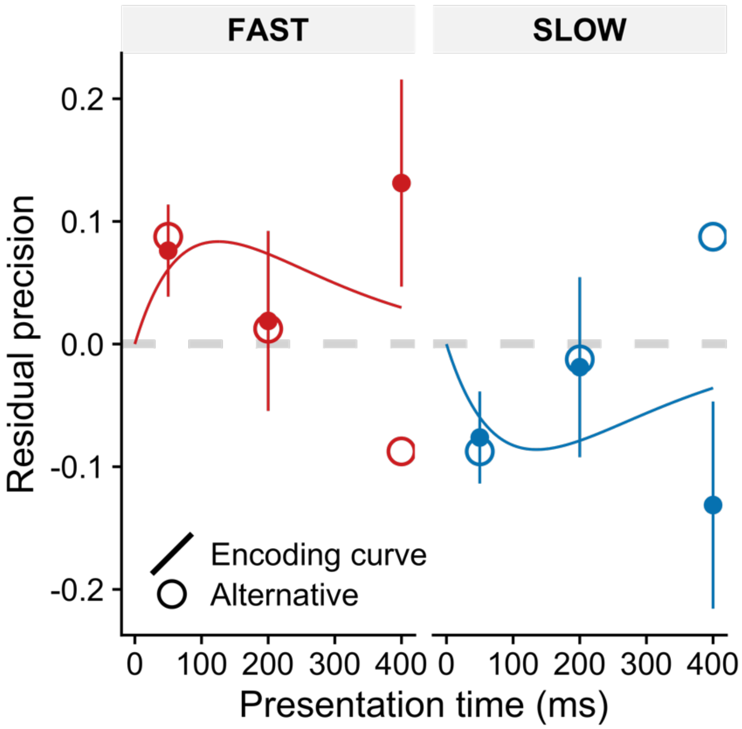


**Supporting Fig. S9.** Testing the alternative explanation that performance benefits and costs scale linearly with the ‘unexpectedness’ of the presentation time (i.e., absolute difference between actual and expected presentation time). Experiment 3: Residual precision (precision – grand average) is plotted at each presentation time for ‘fast’ and ‘slow’ cues. Filled dots with error bars represent data, smooth encoding curve represents encoding speed adaptations and open circles represent alternative explanation. Encoding speed adaptations were fitted for each condition while keeping capacity fixed. The magnitude of benefits and costs were fitted for each condition separately.
